## Supplementary Materials for "Automatic bioinformatic software named entity recognition from literature"

Hao Xuan *et al.*

**This PDF file includes:**

Fig. S1 to S2
Table S1

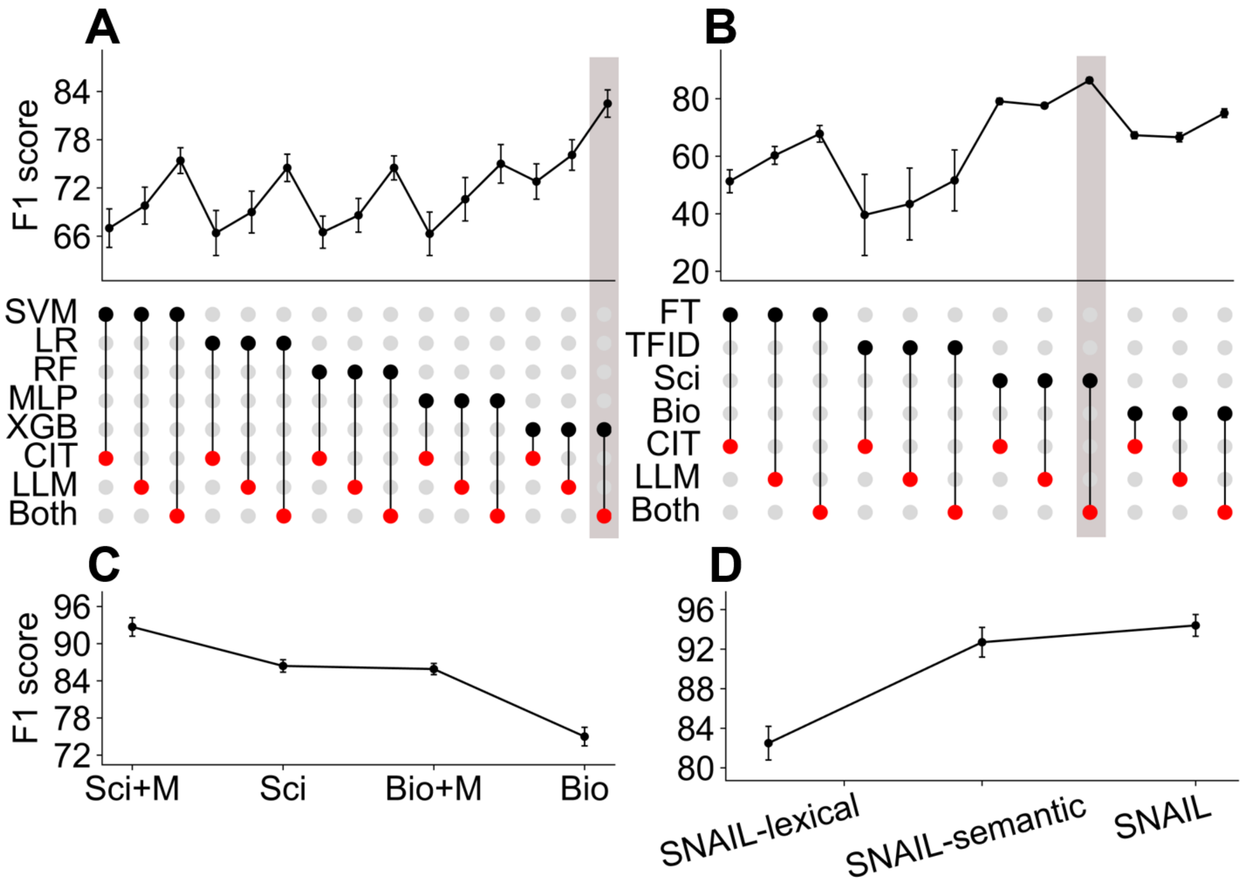


**Fig. S1: Model selection, ablation analysis, and integration of SNAIL components across datasets and training strategies on DS2.** (**A**) Comparison of lexical classifiers for SNAIL-lexical. Models include support vector machine (SVM), logistic regression (LR), random forest (RF), multilayer perceptron (MLP), and extreme gradient boosting (XGBoost, XGB). Black dots indicate the selected lexical classifier, whereas red dots indicate the training dataset configuration: citation-hinted sentences (CIT), large language model–generated sentences (LLM), or the merged dataset containing both sources (Both). The upper panel shows the corresponding F1-scores. The vertical gray shaded bar highlights the optimal configuration. (**B**) Comparison of semantic embedding strategies for SNAIL-semantic using FastText (FT), term frequency–inverse document frequency (TF-IDF, TFID), SciBERT (Sci), and BioBERT (Bio) embeddings coupled with an MLP classifier. Black dots denote the selected embedding model, and red dots denote the training dataset configuration (CIT, LLM, or Both). The upper panel shows the corresponding F1-scores. The vertical gray shaded bar highlights the optimal configuration. (**C**) Effect of explicit masking of the candidate token during semantic model training. Performance is compared for SciBERT (Sci) and BioBERT (Bio) models trained with masking (Sci+M and Bio+M) or without masking (Sci and Bio). (**D**) Performance comparison of the integrated framework components, including SNAIL-lexical, SNAIL-semantic, and the final fused SNAIL model. Error bars represent standard deviation across cross-validation folds.

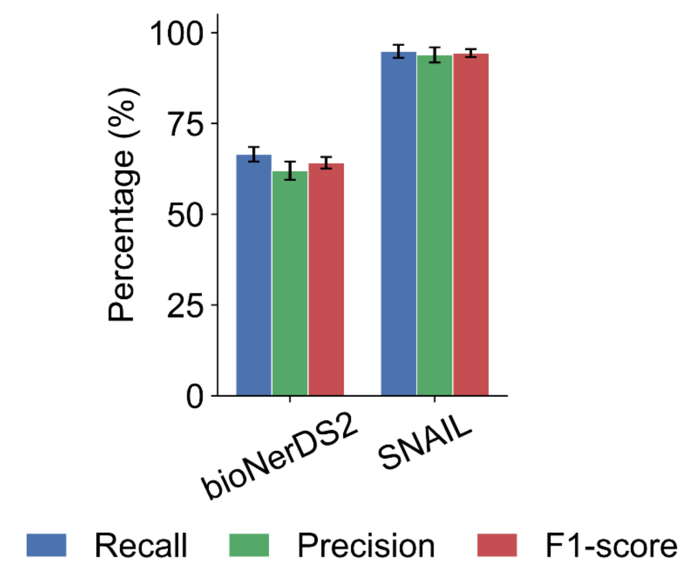


**Fig. S2: Benchmarking of SNAIL against bioNerDS2.** Performance comparison on benchmark datasets. Precision, recall, and F1-score of SNAIL and bioNerDS2 evaluated on DS2. Error bars indicate standard deviation across cross-validation folds.

**Table S1: Generational large language models utilized for benchmark comparisons.** Models are organized by institutional lineage to track the progression from earlier-generation variants to recent state-of-the-art updates. The selected models represent the core evolution of the Claude, Gemini, Grok, and GPT product ecosystems used to evaluate SNAIL performance.

| LLM Ecosystem | Earlier Generation | Latest Generation |
|---|---|---|
| **Claude** | Claude-3.5-Sonnet | Claude-Sonnet-4.6 |
| **Gemini** | Gemini-2.5-Flash | Gemini-3-Flash-preview |
| **Grok** | Grok-4 | Grok-4.20-0309-reasoning |
| **ChatGPT** | GPT-4.1-mini | GPT-5.4-mini |
